# Mapping Histoplasma in Bats and Cave Ecosystems: Evidence from Midwestern Brazil

**DOI:** 10.1101/2025.03.31.641930

**Authors:** João Paulo Romualdo Alarcão Bernardes, Bernardo Guerra Tenório, Joaquim Lucas, Carlos Emilio Molano Paternina, Regianne Kelly Moreira da Silva, Fabián Andrés Hurtado Erazo, Ildinete Silva Pereira, Lucas Gomes de Brito Alves, Paulo Henrique Rosado Arenas, Igor Daniel Bueno-Rocha, Edvard Dias Magalhães, Herdson Renney de Sousa, Hugo Costa Paes, Rosely Maria Zancopé-Oliveira, Daniel Ricardo Matute, Maria Sueli Soares Felipe, Ludmilla Moura de Souza Aguiar, Sébastien Olivier Charneau, André Moraes Nicola, Marcus de Melo Teixeira

**Author notes:** **Correspondence:** Marcus de Melo Teixeira - Núcleo de Medicina Tropical, Faculdade de Medicina, Universidade de Brasília, Campus Universitário Darcy Ribeiro, Asa Norte, Brasília, DF, Brazil CEP 70910-900. First Author and Second Author contributed equally to this work. Author order was determined by Marcus de Melo Teixeira.

## Abstract

Caves serve as natural reservoirs for diverse microbial species due to their unique biotic and abiotic conditions. *Histoplasma* spp. is frequently associated with guano-enriched soil, low luminosity, and high humidity, particularly in Latin America, a region highly endemic for histoplasmosis. Despite the continent’s diverse biomes, local environmental and host distributions of *Histoplasma* remain poorly understood. To address this knowledge gap, we conducted a *Histoplasma*-specific quantitative PCR (qPCR) assay targeting the *hc100* gene on guano samples from seven bat-inhabited caves and tissue samples from 74 bats of nine species in the Federal District of Brazil and surrounding regions. We detected *Histoplasma* DNA in 16 of 80 soil samples (20%) and in 33 bats representing seven species. Among 222 tissue samples (74 lung, 74 spleen, 74 brain), 39 tested positive: 22 lung, 10 spleen, and 7 brain samples. Four bats had *Histoplasma* DNA in both lung and brain, and two in both lung and spleen. By mapping the presence of *Histoplasma* across sampled caves, we identified environmental hotspots of fungal prevalence, emphasizing the need for targeted surveillance.

**Importance:** Our study provides critical insights into the environmental and host distribution of *Histoplasma* spp. in Brazil, identifying caves with high fungal prevalence and demonstrating its presence in multiple bat species. These findings underscore the necessity of public health interventions to mitigate the risk of histoplasmosis among cave visitors in the region. Additionally, we highlight the utility of qPCR for detecting *Histoplasma* in environmental and biological samples, supporting future epidemiological research in Latin America.

## 1. INTRODUCTION

Natural caves are biodiverse ecosystems, having served as habitats for a myriad of species (1). From low to no luminosity, high humidity, constant temperature and high concentrations of organic and inorganic matter, these environments combine specific conditions for the coexistence of cavernicolous species encompassing all five kingdoms (2). From a public health perspective, they may harbor medically relevant species, including viral, bacterial and fungal pathogens (3–5). Among the many microorganisms found in cave environments, *Histoplasma* spp. stands out as a dimorphic fungal genus responsible for histoplasmosis, a systemic mycosis with global distribution (6, 7). In the soil, *Histoplasma* grows as mycelia and produces infectious spores, referred to as micro- and macroconidia. When *Histoplasma*-contaminated soil is disturbed, humans and other mammals can inhale the fungal propagules. Once in the lungs, mammal body temperature (37°C and upwards) cues the transition of these propagules which convert into yeast forms, potentially causing a primary lung infection. (8). The fungus naturally infects bats and is consistently identified through microbiological, serological, or molecular assays across Latin America (9, 10). Bat excrement of these animals is rich in nitrogen and phosphorus. Combined with an acidic pH, mild temperatures, and high humidity, these nutrients create a hospitable environment for *Histoplasma* growth in bat-inhabited caves (11). This tight co-existence suggest that bats might play an important role in the ecology and evolution of *Histoplasma* (10, 12).

Cave visitation has long been associated with histoplasmosis outbreaks (13–16). Brazil has 23,378 catalogued caves, with numerous others awaiting exploration. Of these, 12,279 are estimated to be in the states of Goiás, Minas Gerais and the Federal District (the latter harboring the country’s capital, Brasília), due to regional geological features (17). Cave visitation by speleologists and tourists is an important scientific and ecotourism activity in the midwestern region of Brazil. However, it presents a potential risk to its practitioners due to the high risk of exposure to *Histoplasma* sp. Notably, there have been continuous reports of outbreaks and isolated cases of histoplasmosis in and around the Federal District, potentially associated with cave visits (18–21). However, little is known about the presence and prevalence of *Histoplasma* in their hosts and within cavernicolous environments in those areas. Field studies are highly needed to identify and prevent exposure to *Histoplasma* sp. In this piece, we bridge that gap. We used qPCR-based assays to detect *Histoplasma* in cave environments and bats in the Federal District and surrounding areas. By identifying the presence of *Histoplasma* in caves and hosts, researchers can pinpoint histoplasmosis hotspots, which could inform public policies to reduce exposure for cave visitors.

## 2. MATERIALS AND METHODS

### 2.1 Collection of Soil Samples and DNA extraction

We collected guano samples from eight different caves located in the Federal District and surrounding areas, including Goiás and Minas Gerais states. The locations of these caves are listed in Table 1. Since the caves contained large amounts of organic material, ranging from bat carcasses to guano, as well as visible fungal mycelia (Figure 1), all team members used appropriate personal protective equipment (PPE), including gloves, N95 masks, and protective suits. All sampling procedures were conducted with the support of cave experts and military firefighters, ensuring strict adherence to biosafety protocols. One cave, Gruta do Sal/Fenda II, which has two entrances, was sampled twice in different sections, one year apart, and the collections were analyzed separately. We collected guano samples using disposable spatulas and placed them in labeled sterile 50-mL conical tubes (Fisher Scientific). We placed all samples in Styrofoam boxes at room temperature before DNA extraction. At the end of each collection, we autoclaved and discarded all PPE and sampling tools.

**FIGURE 1.**
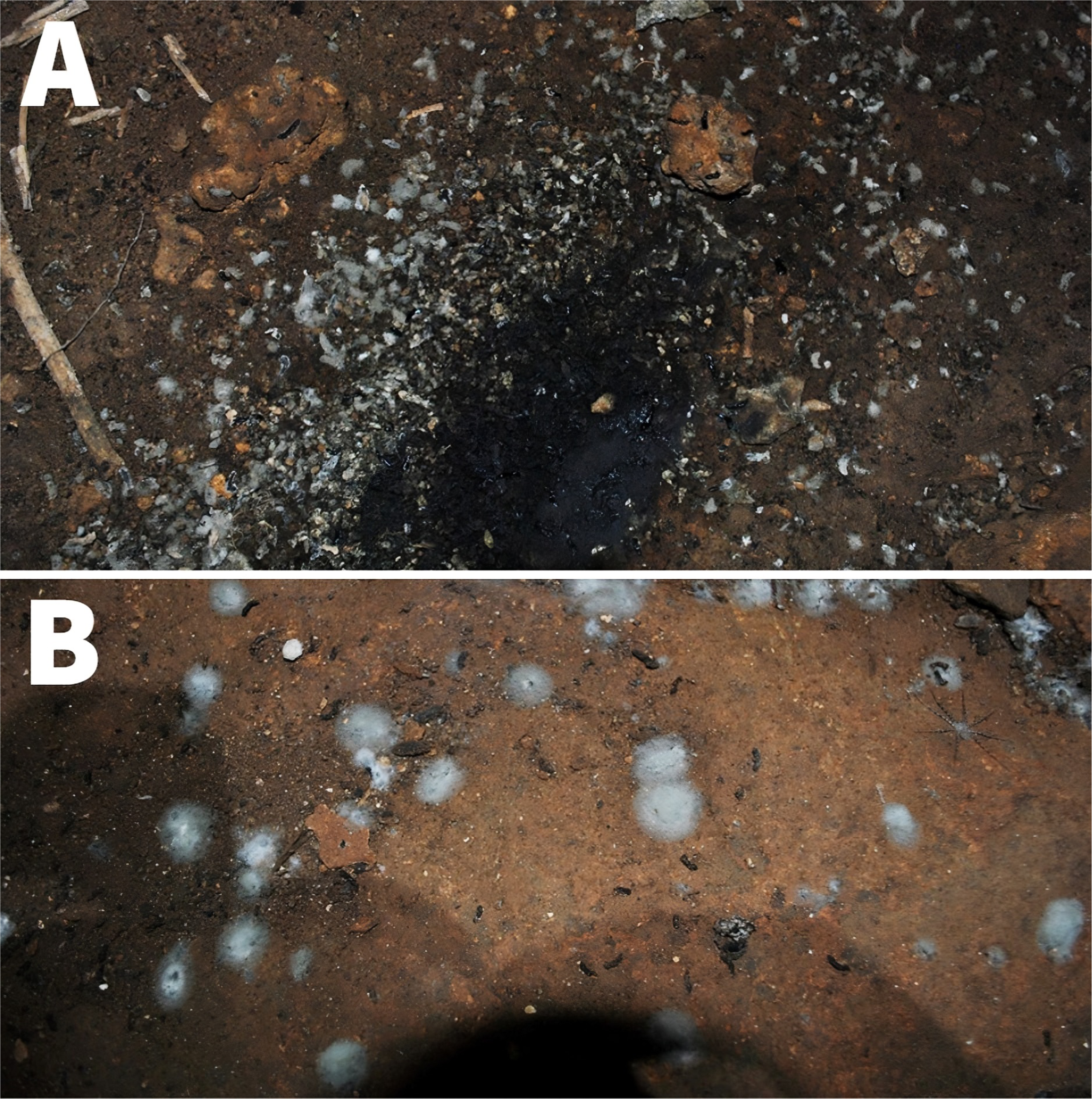
Photographs of chosen sites of collection. (A) Samples were collected on the periphery of the guano puddles while avoiding unnecessary guano overload from the darkest portion. (B) A clearer visualization of mycelia growing from bat guano (white cottonous spots).

**TABLE 1.**
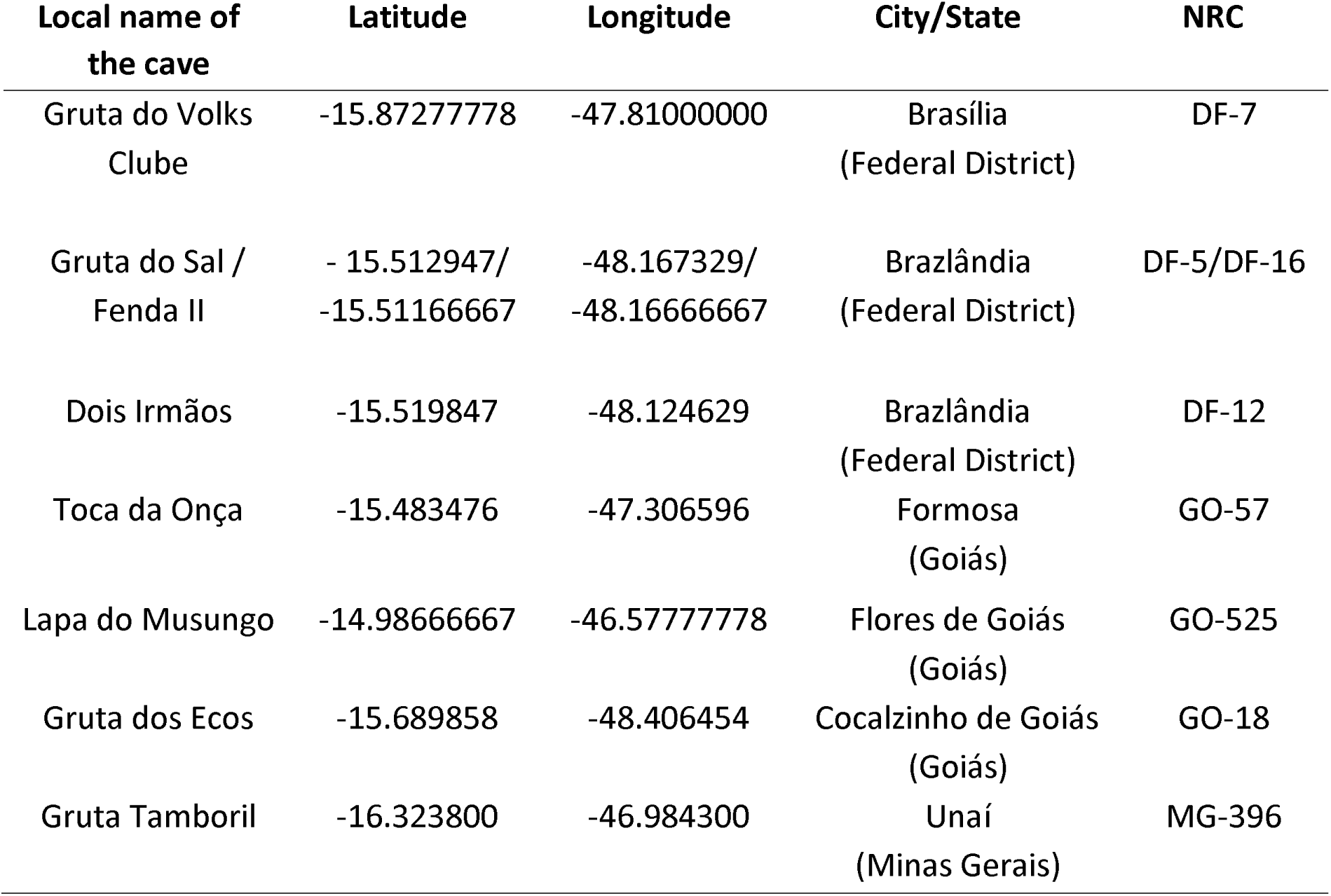
Location of caves sampled for *Histoplasma* sp. detection in Brazil. The NRC (“Cave Registry Number”) is a cave identifier created by a local speleology group (SBE – Brazilian Society of Speleology - https://sbecnc.org.br/). *Gruta* is the Portuguese word for “grotto”.

To perform DNA extraction, we used 0.25 to 0.5 g of soil-containing guano as input material in a class II A2 biosafety cabinet using the DNeasy PowerSoil Kit (QIAGEN). We followed the manufacturer’s protocol but included ten cycles of one minute at 6,500 rpm for the cell lysis step by using a Precellys® Tissue Homogenizer (Bertin Technologies). We quantified the yield of DNA extraction on a Qubit® and on a Nanodrop 2000® instrument (both by ThermoFisher Scientific). All samples were adjusted to a final concentration of 10 ng/μL with nuclease-free water.

### 2.2 Bat capture and DNA extraction

To assess the presence of *Histoplasma* in bats, we captured individuals using mist nets deployed in both urban and rural areas, as well as at cave exits in the Federal District and surrounding regions. We collected bats in 13 different locations in urban, peri-urban and rural areas of the Federal District and its surroundings. At least four mist nets (12 × 3 m) were mounted at each sampling site. The nets stayed open for eight hours starting at sunset (from 6:00 pm to 2:00 am) and we checked for animals every 15 minutes. For this study, we included only adult individuals. We removed captured bats from the net, and placed them in cotton bags except for pregnant or lactating females, which were handled more quickly, banded, and immediately released. All specimens received a unique identification number associated to the collection metadata (capture location, forearm length, weight, age, sex and species). Collected individuals were then transported to the Bat Biology and Conservation Centre at the University of Brasília for euthanasia and morphological characterization. Euthanasia was carried out using isoflurane anesthesia followed by cardiac puncture, in accordance with the protocols previously outlined (22). Dissected organs were stored in RNAlater® (Thermo Fisher) and preserved at -80 °C at the Institute of Biological Sciences of the University of Brasília for subsequent DNA extraction. After all these procedures, the skulls and skins of individuals in good conditions were stored at the Mammal Collection of the Department of Zoology at the University of Brasília. We sought a minimum of 70 adult bats with a balanced sex ratio, evenly distributed across species, including frugivores, insectivores, and hematophagous bats (See Results). All collections were approached by the University of Brasília Institutional Animal Care and Use Committee, process 23106.133703/2020-92, and the Biodiversity Permits and Information System (SISBIO), under protocol number 77229.

Next, DNA extraction from lungs, spleen and brain were performed using the Allprep® DNA/RNA/Protein mini kit (QIAGEN). We followed the manufacturer’s protocol but included three cycles of one minute at 6,500 rpm for the cell lysis step in a Precellys® Tissue Homogenizer (Bertin Technologies) to ensure fungal cell wall disruption. The integrity of DNA was assessed through agarose gel electrophoresis and it was subsequently quantified using spectrometry on a NanoDrop 2000®. DNA samples were adjusted to a final concentration of 10 ng/μL with nuclease-free water.

### 2.3 Validation of the positive control and the qPCR assay

As the positive control, we utilized purified DNA from the strain, CAO4, which belongs to the *Histoplasma* RJ lineage (23). This strain was recovered from a dog assisted at the Evandro Chagas National Institute of Infectious Diseases (INI/Fiocruz) in Rio de Janeiro (23, 24). To validate the positive control for quality, we performed a conventional PCR assay of the universal fungal barcodes targeting the rDNA locus ITS1-5.8S-ITS2 (25). In addition, the positive control sample was subjected to a PCR targeting the gene encoding the hc100 antigen of *Histoplasma* sp. using the 84R22 (5’ GCAGARAATACCYCAAGCC 3’) forward primer and the 15F23 (5’ GTATCCCACAGCATCMYGGAGGT 3’) reverse primer (26). Both conventional PCR reactions were set up in a final volume of 25 µL, containing 12.5 μL of the GoTaq® qPCR Master Mix (Promega Corporation), 0.15 µM of each primer, and 10 ng of template DNA. Thermocycling was performed in a Veriti™ 96-Well Fast Thermal Cycler (Thermo Fisher) with the following conditions: initial denaturation at 95 °C for two minutes; 35 cycles of denaturation at 94 °C for 30 seconds, annealing for 45 seconds, at 55 °C for the ITS locus and 58 °C for the *hc100* locus, and extension at 72 °C for 30 seconds; and a final extension at 72 °C for 5 minutes. PCR products were loaded onto a 1% agarose gel stained with ethidium bromide and submitted to electrophoresis. Gels were documented using a ChemiDoc™ XRS+ imaging system (Bio-Rad).

Finally, we validated the qPCR assay targeting the *hc100* gene using the primers described above and the fluorescent TaqMan probe 45L23 (5’–/56-FAM/CCTTCTTGCAACTYCCYGCGTCT/3BHQ_1/–3’) on the purified DNA of the CAO4 strain. Reactions were performed in a final volume of 20 µL, consisting of 10 µL of the Prime-Time® Gene Expression Master Mix 2X (Integrated DNA Technologies), 0.15 µM of each primer and the probe, and template DNA at concentrations of 0.1 ng, 1 ng, 10 ng, and 100 ng to determine the minimal detection threshold. The qPCR assays were set up in 96-well plates and performed on a 7500 Fast Real-Time PCR System (Thermo Fisher) with the following conditions: initial denaturation at 95 °C for 10 min; plus 50 cycles of denaturation at 94 °C for 15 seconds, annealing at 58 °C for 30 seconds, and extension at 72 °C for 10 seconds. The reaction signal emitted by the TaqMan probe fell within the FAM-TAMRA range. The output data was processed and analyzed using the Design and Analysis 2.6.0 Real-Time PCR System software (Thermo Fisher Scientific). The qPCR experiments aimed at detecting *Histoplasma* sp. in both environmental samples and DNA from bats followed the previously described parameters. These parameters included the use of 10 ng of template DNA and were conducted in triplicate for each sample.

### 2.4 Georeferencing of *Histoplasma* sp. in the Federal District and surrounding areas

Next, we plotted the abundance of qPCR positive assays in a map. The latitude and longitude coordinates of the caves were retrieved from the National Cave Registry of the Brazilian Society of Speleology (NCR/SBE). The cartographic database was taken from the database of the Brazilian Institute for Geography and Statistics (IBGE - https://www.ibge.gov.br/geociencias/organizacao-do-territorio/malhas-territoriais/15774-malhas.html). We used the QGIS Standalone software version 3.22 to combine these datasets and plot the maps.

## 3. RESULTS

### 3.1 Molecular detection of *Histoplasma* in cave environments

We collected 80 soil samples across seven caves located in the Federal District and the states of Goiás and Minas Gerais, as shown in Figure 2. PCR assays targeting both *ITS* and *hc100* successfully amplified DNA fragments of the expected sizes (391 bp and 210 bp, respectively) for the *H. suramericanum* (CAO4) positive control (data not shown). The qPCR assay demonstrated a high sensitivity for the positive control in DNA concentrations ranging from 0.1ng to 100 ng (Figure S1). After standardizing the qPCR assay, we observed that 16 soil samples (20%) were positive for the fungal DNA, and 64 (80%) were negative (Table 2, Figure 2). The 16 positive samples were identified in six of the eight caves sampled as follows: Gruta do Sal Fenda II (2020), 21% positivity (three out of 14 samples); Gruta do Sal / Fenda II (2021), 60% positivity (three out of five); Gruta Dois Irmãos, 11% positivity (three out of 27); Caverna dos Ecos, 50% positivity (three out of six); Toca da Onça, 43% positivity (three out of seven); Gruta do Tamboril, 11% positivity (one out of nine). In contrast, all samples from Gruta do Volks Clube (six samples) and Lapa do Musungo (three samples) were negative for *Histoplasma* sp. (Table 2, Figure 2). The exact location of each cave coupled with positive and negative qPCR results are shown in Figure 2 and Supplementary Table 1.

**FIGURE 2.**
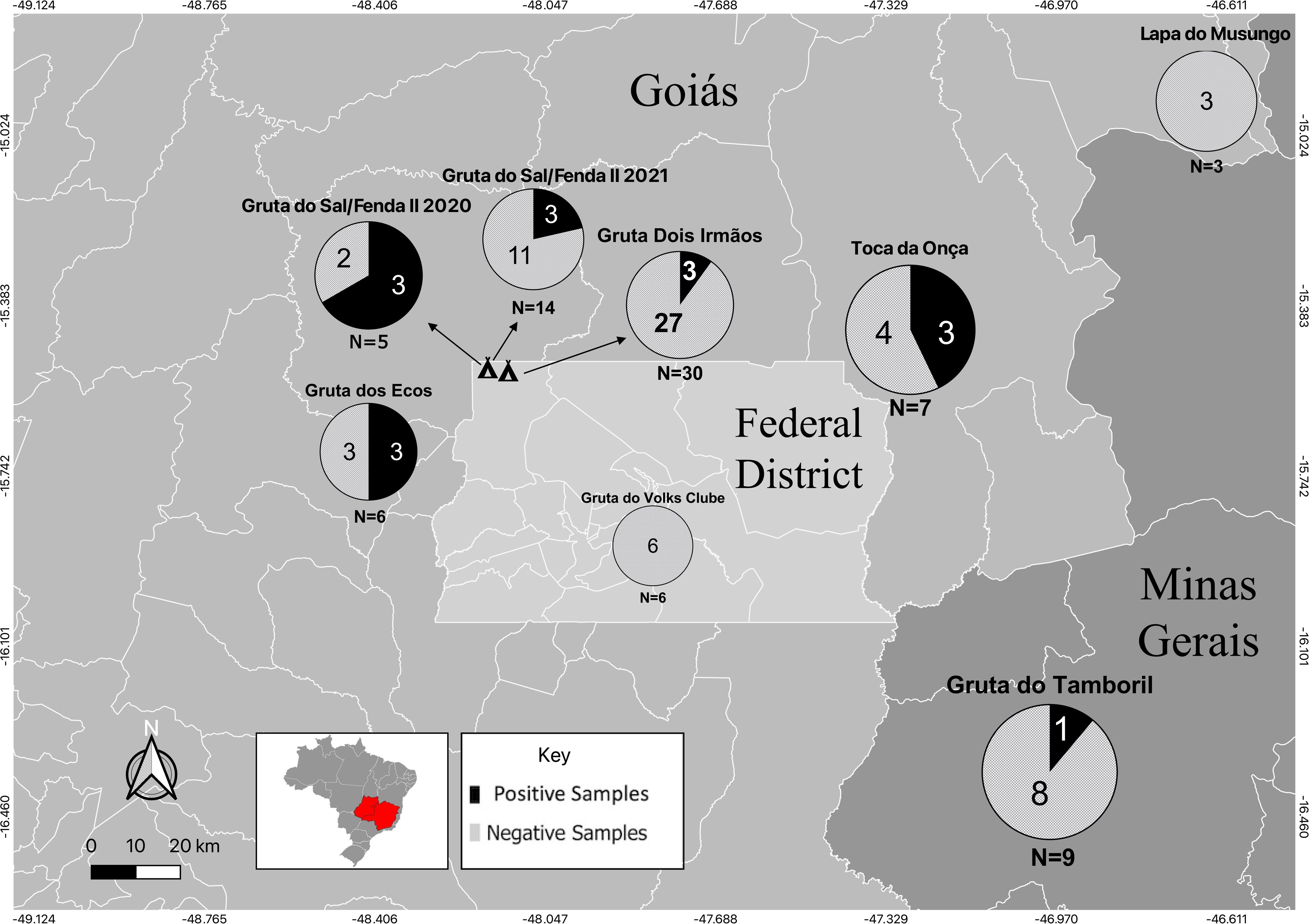
Molecular identification of *Histoplasma* sp. in caves of the Federal District and surrounding areas. The map shows the locations of caves and the results of the qPCR assay for *Histoplasma* sp. in eight different locations. Each circle represents a cave, with the total number of samples (N) indicated below it. Gruta do Sal/Fenda II and Gruta Dois Irmãos, located adjacent to each other, are represented by cave symbols, with arrows indicating their respective results.

**TABLE 2.**
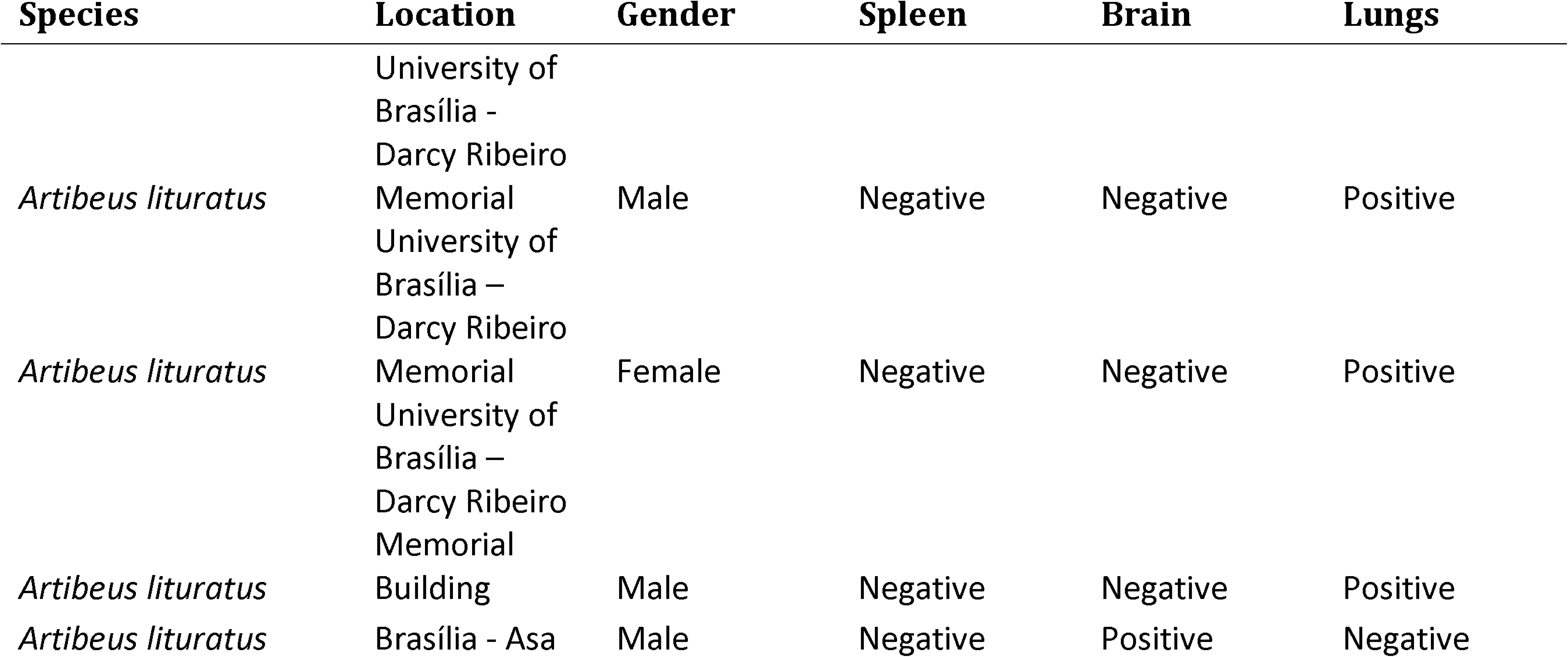
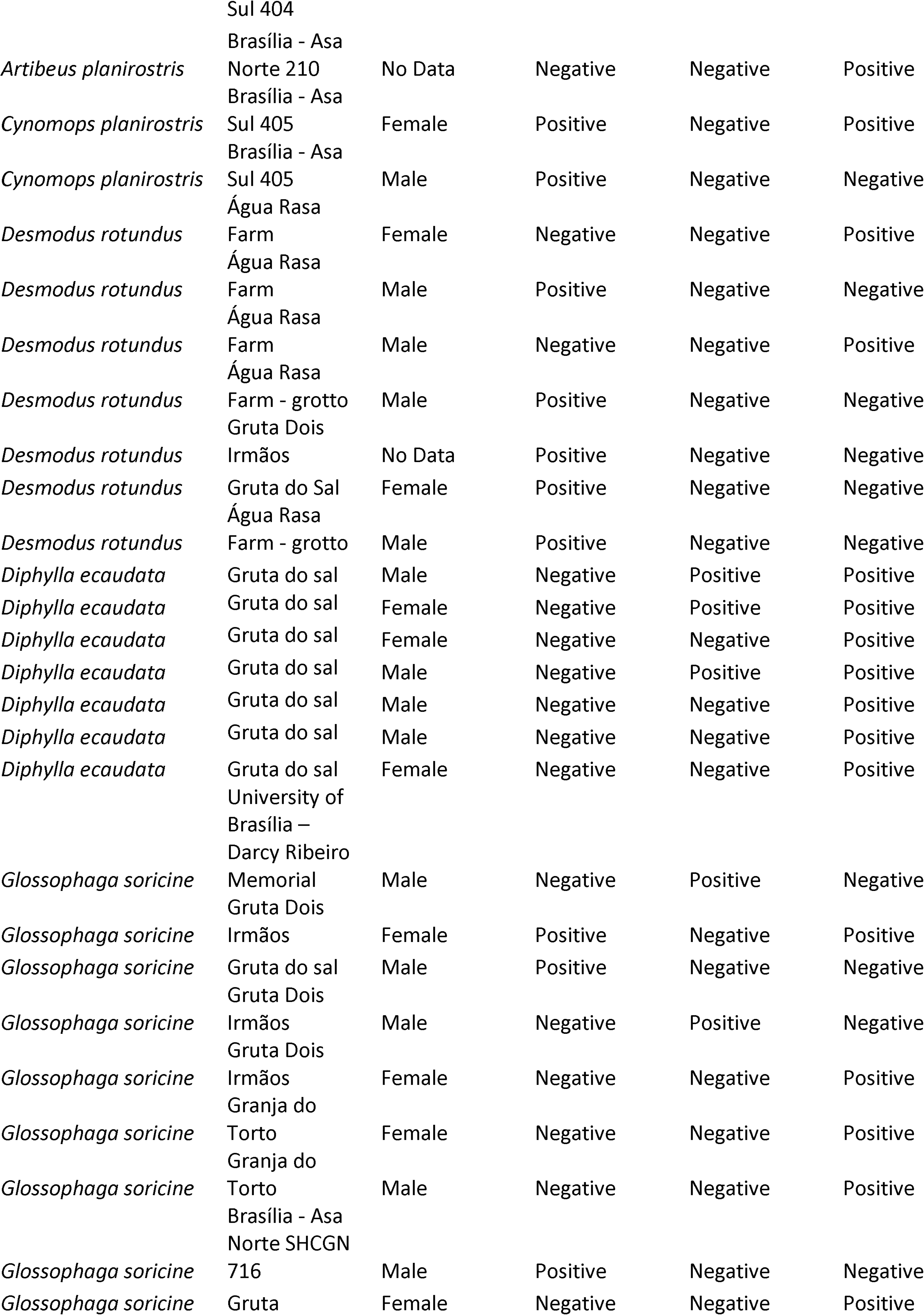
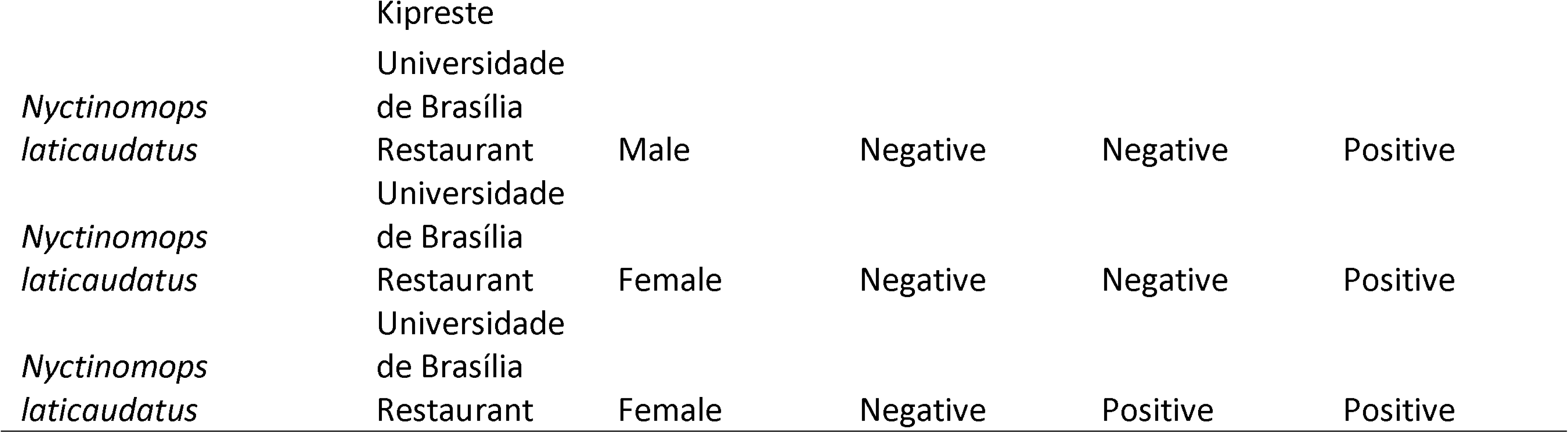
Summary of *hc100* qPCR assays from bats samples positive for *Histoplasma spp*.

### 3.2 Molecular detection of *Histoplasma* in bats

We captured 74 bats, 32 in urban areas and 42 in the rural areas of the Federal District. The sample included 39 males and 32 females, while the sex of three individuals could not be determined. Based on Diaz *et al* . (27) we identified nine species: *Artibeus lituratus* (8), *Artibeus planirostris* (7), *Cynomops planirostris* (5), *Desmodus rotundus* (10), *Diaemus youngi* (10), *Diphylla ecaudata* (10), *Glossophaga soricina* (18), *Molossus molossus* (1) and *Nyctinomops laticaudatus* (5).

*Desmodus rotundus* and *Diphylla ecaudata* , with a prevalence of 70%, were the two species with the highest prevalence of positive tests, followed by *Nyctinomops laticaudatus* (60%), *Artibeus lituratus* and *Glossophaga soricina* (50%), *Cynomops planirostris* (40%), and *A. planirostris* (14%). In contrast, no traces of *Histoplasma* DNA were detected in *Diaemus youngi* and *Molossus molossus*. Among all samples, lung tissue had the highest positivity rate, while brain tissue had the lowest. These results are summarized in Figure 3, Table 2 and Supplementary Table2.

**FIGURE 3.**
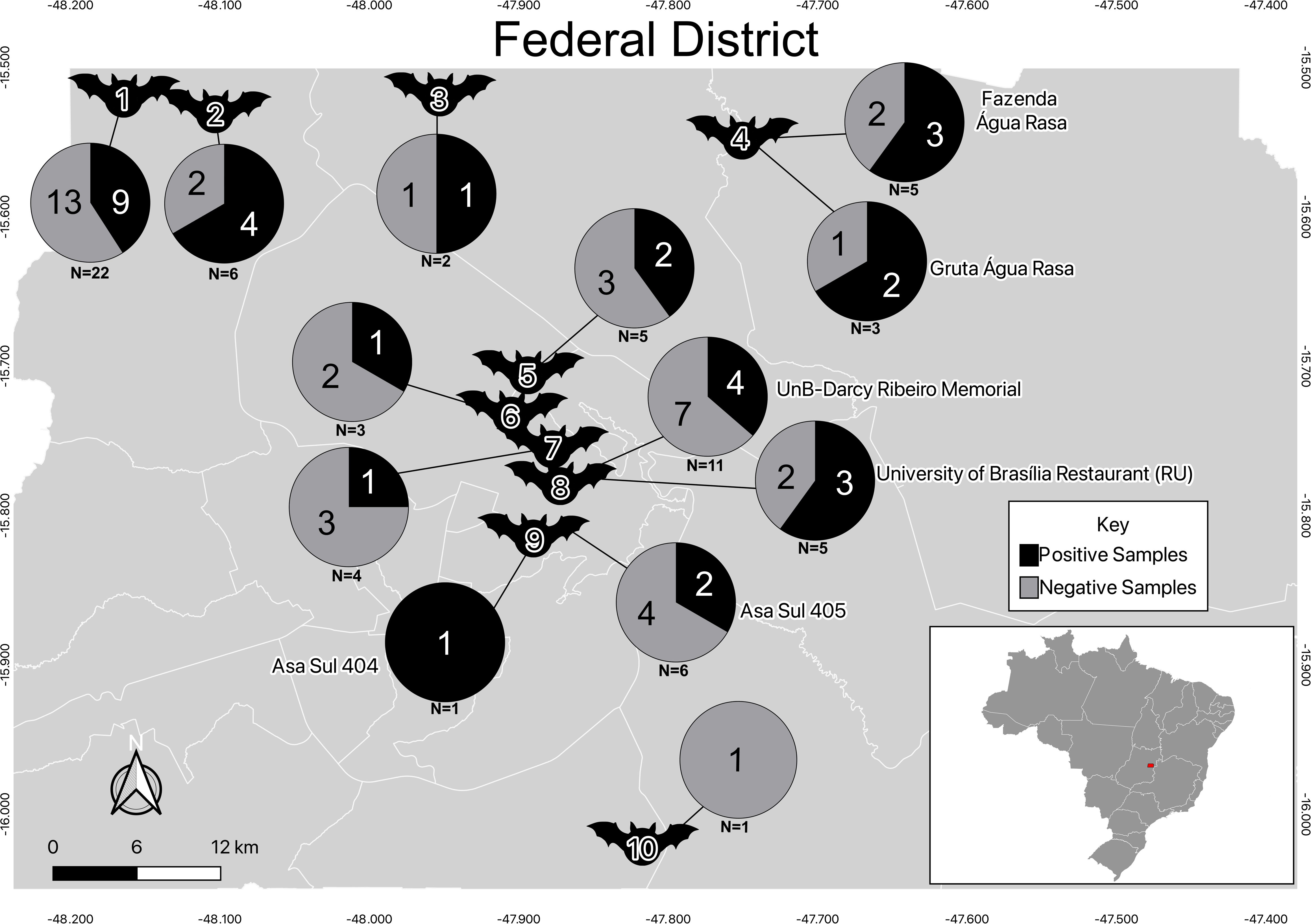
Molecular identification of *Histoplasma sp*. in bats captured in the Federal District. The map illustrates the geographical distribution of bats captured in the Federal District, as well as the results of the qPCR assays. Each bat represents the coordinates of the , Sucapture location, as follows: (1) Gruta do Sal; (2) Gruta Dois Irmãos; (3) Gruta Kipreste; (4) Água Rasa Farm and the grotto in it; (5) Granja do Torto; (6) Asa Norte, SHCGN 716; (7) Asa Norte, SQN 210; (8) University Restaurant and the Darcy Ribeiro Memorial at the University of Brasilia; (9) Asa Sul, SQS 404/405; (10) Jataí Farm. The respective qPCR assay results for each location are depicted in pie charts containing the number of positive and negative samples. The total sample number (N) is displayed under the pie chart for each location.

Male and female bats had similar infection rates, with *Histoplasma* sp. detected in 18 out of 39 (46.1%) males, 13 out of 32 (40.6%) females, and two of the three bats with undetermined sex tested positive for Histoplasma sp. The proportion of infected individuals did not differ among sexes (2-sample test for equality of proportions with continuity correction; X^2^ = 0.0515, df = 1, p-value = 0.821).

Next, we studied whether Histoplasma showed organ tropism in our bat sample (Figure 4). No bat showed infection in all three tissues. Sixteen bats showed positive results only in the lungs, three only in the brain, and eight exclusively in the spleen. Additionally, four bats had positive results in both the lungs and brain, while two tested positive in both the lungs and spleen.

**Figure 4.**
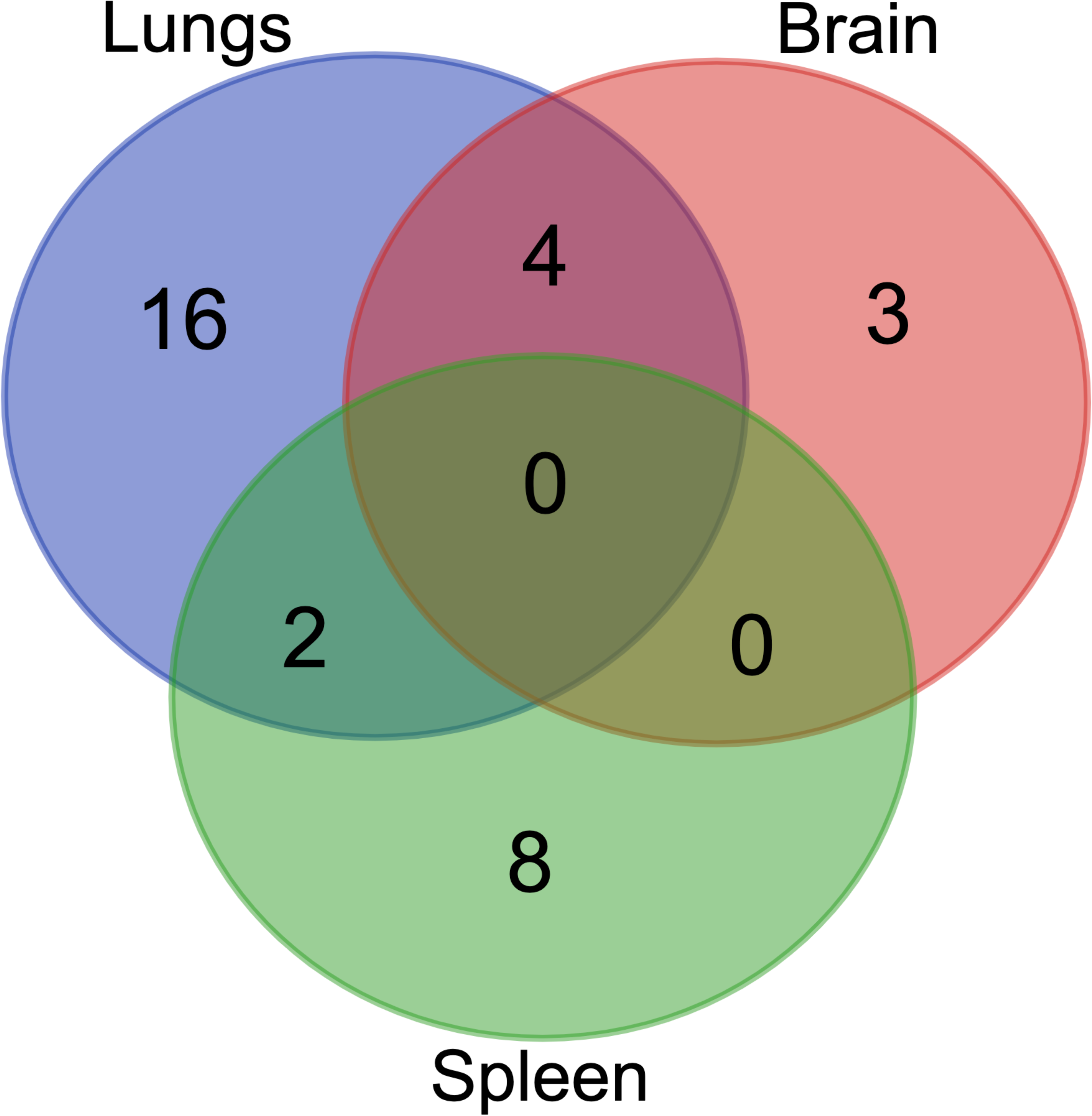
Distribution of Positive Results Across Different Organs in Bats. Venn diagram representing the distribution of positive results across different organs in bats. It also shows the overlap of positive results in the lungs, brain and spleen.

## 4. DISCUSSION

Natural caves provide unique environmental conditions that support diverse ecosystems, including pathogenic fungi such as *Histoplasma* spp. (28). This study investigated the presence of *Histoplasma* in soil samples from seven Brazilian caves in Goiás, Minas Gerais, and the Federal District, regions with numerous caves (17). We also examined bats in the Federal District, recognizing their role as natural reservoirs of the fungus. Our research was prompted by two histoplasmosis outbreaks in a rural area of the Federal District and a high incidence of cases among HIV/AIDS patients (18, 21) underscoring the need to understand mammalian hosts and their environments in fungal transmission. Our results indicate that *Histoplasma* is endemic and occurs in the environment in the Federal District and surrounding areas of Goiás and Minas Gerais (7). Our findings have important implications for detecting *Histoplasma* in environmental and clinical samples, expanding our understanding of its distribution in Brazil, and assessing its epidemiological impact on public health and wildlife reservoirs.

First, our assays demonstrate that a qPCR assay targeting the *hc100* gene is an efficient and reliable tool for detecting *Histoplasma* in environmental and animal samples (Figure 2-3). This method has been validated for detecting *Histoplasma* in chicken manure, cave soil, bird excrement and bat guano, and organic fertilizers (26, 29). Additionally, the assay has been standardized for detecting *Histoplasma* in murine tissues with varying fungal loads (26). Given its high sensitivity, *hc100*-based qPCR has potential for routine clinical diagnosis, though further validation is necessary. Traditional diagnostic methods, such as fungal culture and biopsy, have low sensitivity and require prolonged incubation, necessitating faster molecular alternatives. Moreover, these methods demand biosafety level 3 (BSL-3) laboratories and specialized personnel, emphasizing the need for rapid molecular and immunological alternatives. Comparative studies have shown that PCR performs well in detecting *Histoplasma* DNA in clinical samples (26, 29). Future research should further assess qPCR applicability across different clinical sample types and different species of *Histoplasma*.

Second, our report constitutes a confirmation that *Histoplasma* is often associated to cavernicolous environments (Figure 2-3). Previous reports documented *Histoplasma* in the Ecos and Tamboril caves. Tamboril Cave has been linked to multiple histoplasmosis outbreaks, though most reports stem from media sources rather than peer-reviewed literature. In 2014, researchers conducting a soil survey developed histoplasmosis within ten days despite using personal protective equipment (PPE) during sample collection (20). This case underscores the difficulty of completely preventing exposure in high-risk environments. The cave has since been repeatedly closed, with warning signs posted at the entrance, but unauthorized access continues due to vandalism and disregard for safety measures (30). These challenges highlight the need for more effective control strategies beyond signage. Similarly, Gruta Dois Irmãos was the site of a recent histoplasmosis out-break among firefighters, reinforcing its status as a disease hotspot (18). In contrast, we did not detect *Histoplasma* in Volks Clube and Lapa Musungo caves, possibly due to sampling limitations or true fungal absence. The metropolitan location of Volks Clube Cave may also inhibit fungal growth due to anthropogenic factors (31). Additional systematic sampling is required to confirm fungal absence.

Our findings also inform our understanding of the relationship between bats and *Histoplasma*. Previous studies have reported that ∼25% of bats in Brazil are infected with *Histoplasma* (32). Bats from Guyana show a similar infection rate (33). Wild bats from Argentina show (90%) a higher rate of infection. Different samplings from Mexico show a wide range of infection rates (30-95%) which might reflect variance in community assembly across geography (33). This study confirmed *Histoplasma* DNA in the lungs, spleen, and brain of seven bat species: *Artibeus lituratus* , *A. planirostris* , *Cynomops planirostris* , *Desmodus rotundus* , *Diphylla ecaudata* , *Glossophaga soricina* , and *Nyctinomops laticaudatus*. Previous research has identified *Artibeus lituratus* , *Desmodus rotundus* , *Glossophaga soricina* and *Nyctinomops laticaudatus* species as carriers of *Histoplasma* infections (34). Thus, we provide valuable data from *A. planirostris*, *Cynomops planirostris* and *Diphylla ecaudata* species which have never been previously described as natural hosts for *Histoplasma*. (34). Two species, *Diaemus youngi* and *Molossus molossus*, tested negative. The former has not been reported for *Histoplasma* but the latter has in numerous occasions (9, 34). This may reflect sample size limitations, as only one *M. molossus* individual was analyzed, or potential resistance in *D. youngi*. The phylogenetic branch of fungal infections, and specifically of *Histoplasma*, in bats remains an open area of research.

Our study is also the first to detect *Histoplasma* DNA in the brain of bats, raising questions about potential central nervous system (CNS) involvement (Table 2). Despite the association between bats and *Histoplasma*, the clinical manifestations (if any) of histoplasmosis in bats remains largely unknown. While the presence of fungal DNA does not confirm active infection, it suggests that *Histoplasma* may occasionally reach the CNS. This finding warrants further investigation into fungal dissemination pathways in bats. In humans, CNS histoplasmosis is rare but can occur in immunocompromised individuals, leading to severe complications such as meningitis or encephalitis (35). Our results highlight the need for future research to determine whether *Histoplasma* can cause neurological disease in bats and assess its implications for transmission dynamics.

## 5. CONCLUSIONS

Our study confirmed the presence of *Histoplasma* in caves from Goiás, Minas Gerais, and the Federal District, as well as in bats from both urban and rural areas of the Brazilian capital. Notably, some of these caves are open to researchers and the public without any safety warnings or protective measures to prevent fungal exposure. Our findings contribute to ongoing efforts to monitor microbial activity in caves. Multiple studies have frequently detected *Histoplasma* in caves across endemic regions, highlighting their role as hotspots for fungal transmission. Additionally, the long-distance migration of bats increases the risk of *Histoplasma* spreading to new areas, expanding the geographic range of histoplasmosis out-breaks (9, 36). To mitigate public health risks, we recommend environmental education initiatives targeting local communities, ecotourism operators, and first responders. Training pro-grams should emphasize histoplasmosis prevention and exposure reduction, particularly in regions with confirmed fungal presence. Regular soil testing during both wet and dry seasons is essential to assess fluctuations in *Histoplasma* distribution. Seasonal monitoring will enhance risk assessment by identifying environmental conditions that promote fungal proliferation. These efforts can guide evidence-based policies for cave management and visitor safety.

## ACKNOWLEDGMENTS

DRM was supported by the National Institute of Allergy and Infectious Diseases (NIAID) of the National Institutes of Health (NIH) under Award R01AI153523. M.M.T was supported by Fundação de Apoio à Pesquisa do Distrito Federal (FAP-DF) grant number 00193-00001871/2023-99 and Conselho Nacional de Desenvolvimento Científico e Tecnológico (CNPq) grant number 314119/2023-0.

## Notes

### Competing Interest Statement

The authors have declared no competing interest.

